# Modulation of electrocorticographic seizure features predicts response to closed-loop brain stimulation

**DOI:** 10.1101/408567

**Authors:** Vasileios Kokkinos, Nathaniel D. Sisterson, Thomas A. Wozny, R. Mark Richardson

## Abstract

Why does closed-loop invasive brain stimulation improve seizure control in some patients with epilepsy, but not others? The RNS System, the only FDA-approved bi-directional brain-computer interface, has been shown to improve seizure control in patients with refractory epilepsy, although the mechanisms behind this success are undefined. We analyzed recordings from the RNS System and discovered two main categories of electrophysiological signatures of stimulation-induced modulation of the seizure network. Direct effects included ictal inhibition and early frequency modulation but did not correlate with improved clinical outcomes. Only indirect effects, those occurring remote from triggered stimulation, predicted improved clinical outcomes. These indirect effects, which included spontaneous ictal inhibition, frequency modulation, fragmentation, and ictal duration modulation, may reflect progressive local epileptogenic network compartmentalization that hinders the spread of pathological synchrony from recruiting neuronal populations. Our findings suggest that responsiveness to RNS may be explained by chronic stimulation-induced modulation of seizure network activity, rather than direct effects on each detected seizure.

## Introduction

How does closed-loop neurostimulation reduce seizure frequency in patients with epilepsy? Modulation of seizure activity by applying electrical current directly to the cortex was reported by Penfield as early as 1945 (Penfield and Jasper, 1954). Sixty years later, a responsive neurostimulator was developed to automatically analyze electrocortical potentials in order to detect seizures and rapidly deliver electrical stimulation, with the goal of suppressing seizure activity (Kossof *et al*., 2004). The RNS System is a closed-loop, responsive neurostimulation system that was FDA-approved as an alternative treatment for patients suffering from drug-resistant focal epilepsy, who are not surgical candidates (Stacey and Litt, 2008). The device records brain electrocorticographic (ECoG) activity from four recording channels in two bi-directionally coupled leads that also deliver stimulation.

Cortical electrical stimulation, in an open-loop acute and/or subacute mode of operation, has been shown to suppress electrographic epileptiform discharges (Velasco *et al*., 2000; Yamamoto *et al*., 2002; Kossoff *et al*., 2004; Kinoshita *et al*., 2004, 2005), as well as to reduce seizure frequency (Velasco *et al*., 2000; Yamamoto *et al*, 2006; Elisevich *et al*., 2006; Velasco *et al*., 2009; Child *et al*., 2014; Ludstrom *et al*., 2016, 2017; Valentin *et al*., 2017; Kerezoudis *et al*., 2017). There is Class 1 evidence for the efficacy of RNS in seizure reduction, with a 44% seizure reduction at 1 year post-implantation, 53% at 2 years (Heck *et al*., 2014), and a range of 48-66% between the 3rd and 6th post-implantation years in open-label continuation studies (Bergey *et al*., 2015). At 6 years, a median 70% of patients with both mesio-temporal and neocortical seizure onset experienced significant reduction in seizure frequency; 26-29% benefited with a post-implantation seizure-free period of at least 6 months, and about 15% experienced 1 year or longer without seizures (Geller *et al*., 2017; Jobst *et al*., 2017). Thus, although the RNS System provides improved seizure control and quality of life in pharmaco-resistant epilepsy, its mechanism of action is unknown, and its overall efficacy remains suboptimal. A critical challenge for the fields of epilepsy and neuromodulation, therefore, is to discover therapeutic mechanisms of closed-loop brain stimulation.

Historically, the primary hypothesis for the mechanism of action of responsive neurostimulation has been the direct inhibition of ongoing ictal activity by triggered electrical stimulation (Lesser *et al*., 1999; Kossoff *et al*., 2004; Skarpaas and Morrell, 2009; Morrell and Halpern, 2016). Isolated samples of recordings and corresponding spectrograms that support this hypothesis have been presented sporadically in the literature (Skarpaas and Morrell, 2009; Thomas and Jobst, 2015; Geller et al., 2017; Jobst *et al*., 2017), but no previous studies have undertaken an in depth analysis of the brain’s response to chronic closed-loop stimulation events. We tested the hypothesis that clinical efficacy arises from successful detection-triggered electrical stimulation and subsequent direct termination of seizure activity, but instead found evidence for an altogether different therapeutic mechanism.

## Materials and Methods

### Participants and implantation

All patients participating in this study had a confirmed diagnosis of focal epilepsy according to current ILAE criteria (Berg *et al*., 2010; Fisher *et al*., 2017). Each patient underwent an investigative intracranial recording procedure, either by subdural ECoG, or by robotic-assisted stereotactic EEG (sEEG), to identify the focus and extent of their epileptogenic zone. After a review of all available patient data during weekly multidisciplinary epilepsy conferences and consideration of available therapeutic options including resective and ablative procedures, closed-loop neurostimulation therapy (RNS, NeuroPace, Mountain View, CA, USA) was recommended. Data from all patients implanted with the RNS System between January 2015 and June 2017 were included in this study, which was approved by the University of Pittsburgh Institutional Review Board (IRB).

RNS leads were positioned as closely as possible to the recorded and/or hypothesis-derived epileptogenic regions. Patients with a diagnosis of neocortical epilepsy onset were implanted either with strips placed over the focus (patient 10), or depth leads placed vertically through the focus (patients 1, 5, and 7), or a combination of both (patient 2). Patients with a diagnosis of malformations of cortical development were implanted with depth leads across the posterior-anterior direction of the lesion (patients 8 and 11). Patients with a diagnosis of mesio-temporal epilepsy were implanted with depth electrodes placed across the posterior-anterior axis of the hippocampus (patients 3, 4, 6, and 9) (Figure 1, Supplementary Figure 1). Assessment of electrode locations was performed by fusion of the post-surgical CT with the pre-surgical MRI (EpiNav, UCL, UK) (Nowell *et al*., 2015).

**Figure 1.**
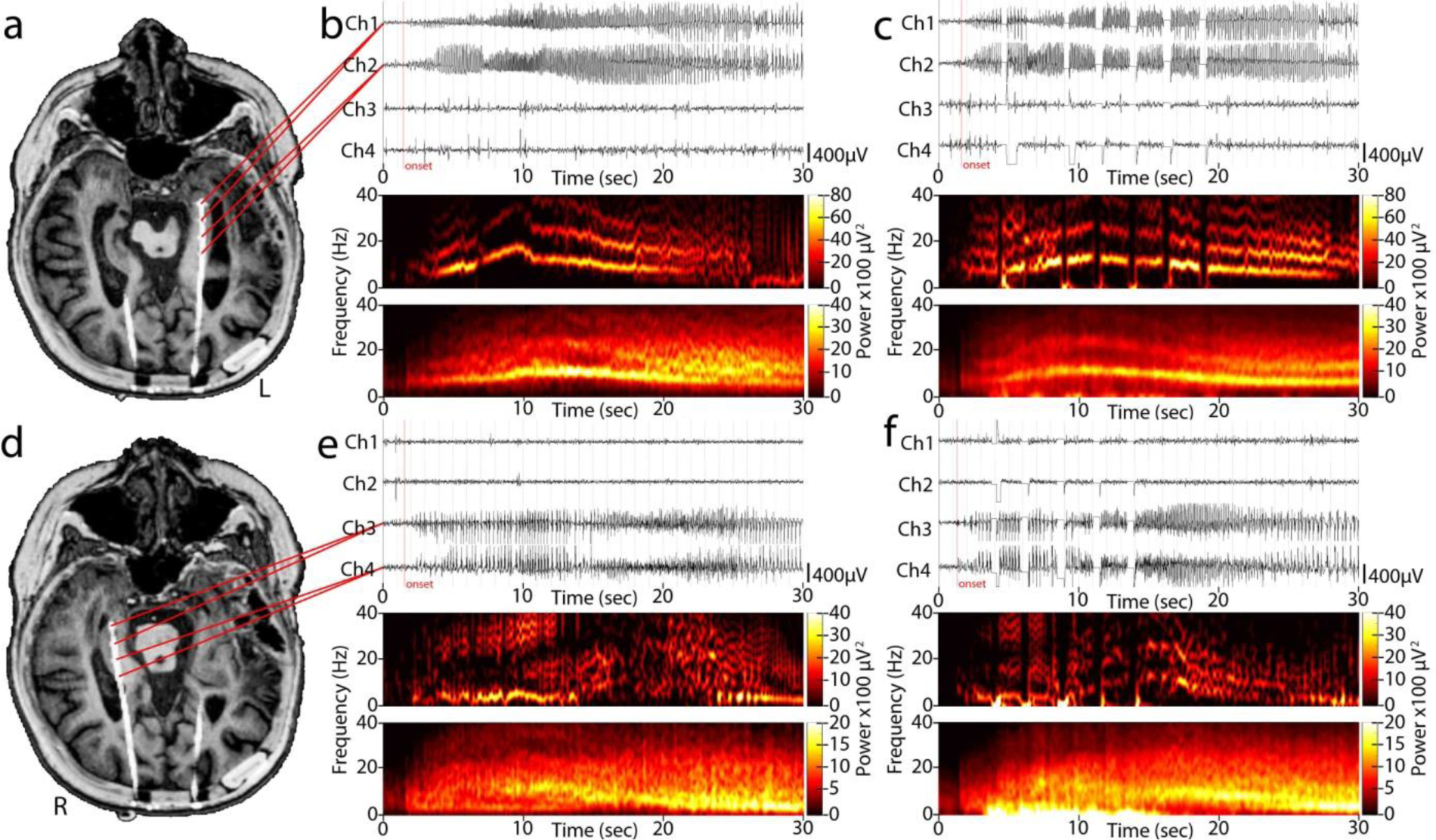
RNS implantation, data processing methodology. (a) Pre-operative MRI and post-implantation CT fused image, aligned in the axial plane across the trajectory of the implanted RNS lead in the left hippocampus of patient 6. (b) The two distal anterior hippocampal contacts provide bipolar channel 1, and the two proximal posterior hippocampal ones form bipolar channel 2 (top), that record a unilateral electrographic seizure pattern in the left hippocampus during the baseline, free of stimulation, period. For channel 2, the lead electrographic seizure patterns onset channel (red line), a 3D time-frequency power matrix is generated, depicting the spectral evolution of the electrographic seizure patterns in time (middle). For all visually confirmed electrographic seizure patterns, the average of all spectrograms is calculated (bottom), aligned in time using the marked onset point as reference. (c) Respective data to (b), for the 1st programming epoch (programming epoch 1) where stimulation was activated. During stimulation the amplifier is disconnected, thereby generating a rectangular pulse artifact in the time domain (top) and a low frequency artifact, accompanied by broadband cancellation, in the frequency domain (middle). Only when stimulation occurs in regular intervals after the electrographic onset, the spectral artifact remains prominent in the averaged spectrogram (bottom). Note the similarity of the spectral traces generated by the prominent oscillating electrographic seizure patterns frequencies between the baseline and stimulation epochs, both in single samples as well as the averaged matrix. Sections (d), (e), and (f) show the respective imaging, data, and processing for an independent right hippocampal electrographic seizure pattern prominent on channel 3 of the same patient.

### Data acquisition

RNS-recorded ECoG data were obtained from NeuroPace upon request via an encrypted mass storage device. Additional RNS metadata, including recording and detection settings, were collected directly from the NeuroPace Patient Data Management System (PDMS) using automated in-house custom-built software (BRAINStim; Sisterson *et al*., under review). All recordings consisted of a 90s duration, 4-channel ECoG that was online band-pass filtered at 4- 125Hz before being sampled at 250Hz and digitized by a 10-bit ADC. ECoG channels were collected using a bipolar recording montage between neighboring electrode contacts (Figure 1, Supplementary Figure 1) with the case of the RNS pulse generator acting as the amplifier ground. All electrode impedances measured below 1kOhm for all recordings.

Both scheduled and detection-triggered ECoG recordings were obtained. Scheduled recordings were triggered by the RNS device’s onboard clock to occur either every 12 or 24 hours and offered a continued sampling of spontaneous neurophysiologic activity. Detection-triggered recordings were initiated by using one of the RNS device’s onboard closed-loop algorithms. Of the three available detector paradigms, the line-length detector was used most frequently followed by the bandpass algorithm; the “area under the curve” detector was not used in this cohort. All patients were instructed to download the RNS raw ECoG data daily to a local computer using a transcutaneous telemetry wand, which was in turn uploaded to the NeuroPace PDMS on a weekly basis.

Immediately post-implantation, the device was set to passive recording mode for approximately 1 month, during which no stimulation was delivered in order to record baseline activity. This recording is referred to as the baseline epoch. Once baseline activity was reviewed, stimulation parameters were configured and activated. In turn, the device delivered detection-triggered stimulation therapy, and parameters were periodically modified in subsequent clinic visits based on evaluation of seizure control status. The mean time from implantation to activation of responsive stimulation was 22 weeks (SD: 8.26). The time interval during which RNS parameters remain unchanged is referred to as a programming epoch.

### Data processing

Electrographic seizure patterns were visually identified by an experienced epilepsy neurophysiologist (V.K.), and their onset was annotated by a cursor marker. The term electrographic seizure pattern is used instead of the term “seizure,” as the device provides no information regarding the manifestation of electrographic events. The electrographic seizure pattern onset was defined as the point in time after which the ECoG recording background was no longer interictal and was followed by a paroxysmal discharge of seizure features and morphology developing over time. Interictal background, both in the awake and sleep states, was appreciated from recorded ECoGs that did not contain electrographic seizure patterns. Electrographic seizure patterns were secondarily clustered in the following groups, those: 1) with a clearly recorded onset versus a preceding interictal background of a minimum 5 s that did not receive stimulation, either because they belong to the baseline period, or because they were not stimulated due to detection failure or exhaustion of the programmed RNS daily stimulation dosage (Supplementary Figure 2a1, b1); 2) with a clearly recorded onset versus a pre-ictal normal background of a minimum 5 s, that received at least one stimulation pulse after the identified onset (Supplementary Figure 2a2, b2); 3) without a recorded onset due to the RNS regularly overwriting its internal memory storage; 4) with a clearly recorded onset versus a pre-seizure normal background of a minimum 5 s, that received at least one stimulation pulse before or during the onset (Supplementary Figure 2a3, b3, a4, b4). Only the first two groups of recorded electrographic seizure patterns were used for evaluation of modulatory stimulation effects.

In order to prepare the data for evaluation in the frequency domain, all electrographic seizure patterns meeting the criteria of the previous paragraph were separated in files of 65 s (starting −5 s before the identified onset, ending 60 s after the onset). Spectral analysis was performed for each sample by means of the Fast Fourier Transform (FFT) of a 128-point window and a 125-point overlap, for frequencies from 0.05 - 60 Hz at a step of 0.05 Hz, using Matlab (The Mathworks, Natic, MA, USA). The power dimension of the spectrum was represented in absolute values and displayed in a linear colored greyscale. For each patient, electrographic seizure patterns were clustered per epoch (i.e. baseline and for each separate programming epoch). In turn, group averages from the individual 3D time-frequency power tables were created for each electrographic seizure pattern cluster per epoch. Homogeneity among clustered items for spectral averaging was assured by the fact that: a) all spectral tables have the same time-frequency dimension size and scale, b) all electrographic seizure pattern onset markers were placed before seizure-like features common across all clustered samples (e.g. a gamma burst, a theta oscillation, or a distinctive slow deflection, etc.), c) clusters comprised of ECoG events whose seizure-like electrographic patterns shared visually appreciable common temporal features, based on the following criteria: morphology, hemispheric distribution, temporal evolution, spread/propagation and duration. In patients with bilateral implantation schemes, independent left and right electrographic seizure patterns, as well as bilateral ones, were marked and grouped separately.

We categorized electrical stimulation effects and clustered the electrographic seizure patterns of each patient as: 1) direct effects, where the recorded events manifested systematic time and/or frequency changes in the immediate post-stimulation interval, and 2) indirect effects, where the recorded events manifested time and/or frequency changes before and/or long after stimulation, and/or could not be attributed to direct stimulation. Changes in the duration of electrographic seizure patterns were scored to be meaningful if they exceeded 25% of the mean duration of the respective averaged baseline patterns. Changes in the spectral content of the electrographic seizure patterns were scored to be meaningful if the they diverged by more than 25% from the respective averaged baseline frequency. Interruptions in the temporal progression of electrographic seizure patterns were scored as discontinuities/fragmentations if there was a return to normal background levels for 0.5 s to 3 s; interruptions of less than 0.5 s were not scored as discontinuities/fragmentations, while those exceeding 3 s were scored as separate electrographic events. Seizure fragmentations of this temporal resolution (0.5 s to 3 s) were described as coarse. Increases in the time interval between consecutive seizure spike discharges were scored as fragmentations if their duration exceeded the respective mean baseline interval by a minimum of 100%. Seizure fragmentations of this temporal resolution (100 ms to 1 s) were described as fine.

### Outcome and statistics

Post-implantation outcomes were derived by extended personal impact of epilepsy scale (PIES) questionnaires (Fisher *et al*., 2015). PIES questionnaires were supplemented with 3 variables of interest subjectively describing seizure manifestation: 1) the mean frequency of seizure occurrence before and after RNS implantation, per month in absolute numbers, 2) the estimated mean severity of seizures, on a scale of 1 to 5 (1: not severe, 5: very severe), and 3) the mean duration of seizures, in minutes. (Table 1).

**Table 1.** Patients, closed-loop stimulation effects, and outcomes (Y: present, X: not present, (-) decrease, (+) increase, R: right, L: left, B: bilateral; Onset weeks refer to the time an effect was first observed after activation of RNS stimulation; Percentages in the Engel scale column represent reported reduction in seizure frequency - SF, severity – SS, and duration – SD, where negative values express increase of variables).

| PATIENTS | AGE | SEX | IMPLANTATION |  | DIRECT STIMULATION EFFECTS |  | INDIRECT STIMULATION EFFECTS |  |  |  |  | OUTCOME (ENGEL SCALE) |
| --- | --- | --- | --- | --- | --- | --- | --- | --- | --- | --- | --- | --- |
|  |  |  | SITES & PATHOLOGY | MONTHS | DIRECT INHIBITION (Onset week) | FREQUENCY MODULATION (Onset week) | SPONTANEOUS ATTENUATION (Onset week) | FREQUENCY MODULATION (Onset week) | COARSE FRAGMENTATION (Onset week) | FINE FRAGMENTATION (Onset week) | MEAN ICTUS DURATION (Onset week) |  |
| 1 | 21 | M | R Fronto-Parietal PMG | 6 | X | X | X | Y (w0) | X | X | X | IIB<br>SF: 98.2%<br>SS: 33.3%<br>SD: 0.0% |
| 2 | 44 | F | L Basal Temporal AVM | 22 | Y (w28) | Y (w21) | X | X | X | X | X | IVB<br>SF: 0.0%<br>SS: 0.0%<br>SD: 0.0% |
| 3 | 24 | F | B Mesial Temporal No lesion | 13 | Y (w20) | X | X | X | X | X | X | IVB<br>SF: 0.0%<br>SS: 0.0%<br>SD: 0.0% |
| 4 | 40 | F | B Mesial Temporal No lesion | 10 | X | Y (w2) | X | X | X | X | X | IVB<br>SF: 0.0%<br>SS: 0.0%<br>SD: 0.0% |
| 5 | 29 | M | R Fronto-Parietal PMG/Heterotopia | 5 | X | Y (w5) | X | Y (w9) | X | X | Y (-) (w3) | IIB<br>SF: 72.3%<br>SS: 8.8%<br>SD: 20.0% |
| 6 | 24 | F | B Mesial Temporal No lesion | 15 | X | X | X | X | X | Y (+) (w60) | X | IVA<br>SF: 0.0%<br>SS: 26.3%<br>SD: 15.7% |
| 7 | 36 | F | R Mesial Frontal No lesion | 19 | X | X | X | Y (w1) | X | X | Y (-) (w12) | IB<br>SF: 100.0%<br>SS: 100.0%<br>SD: 100.0% |
| 8 | 40 | F | L Fronto-Parietal Heterotopia | 9 | Y (w4) | X | X | Y (w20) | Y (w19) | X | X | IIIA<br>SF: 90.0%<br>SS: 81.8%<br>SD: 92.7% |
| 9 | 32 | M | B Mesial Temporal | 8 | X | X | X | X | X | X | Y (+) (w7) | IVC<br>SF: -11.4% |
|  |  |  | No lesion |  |  |  |  |  |  |  |  | SS: 6.6%<br>SD: 50.0% |
| 10 | 65 | F | B Temporal<br>No lesion | 13 | **Y**<br>(w5) | X | X | X | **Y**<br>(w8) | X | **Y (-)**<br>(w53) | IB<br>SF: 100.0%<br>SS: 100.0%<br>SD: 100.0% |
| 11 | 49 | F | B Occipito-Temporal<br>Heterotopia | 26 | X | X | **Y**<br>(w9) | X | X | X | **Y (-)**<br>(w81) | IIIA<br>SF: 66.7%<br>SS: 77.7%<br>SD: 50.0% |

To describe the association between outcome and the presence of a direct or indirect modulation effect, we first grouped patients as either responders (Engel class III or better) or non-responders (Engel class IV), based on the scores of the 3 seizure manifestation variables (Engel *et al*., 1993). Next, we computed two binary variables representing the presence or absence of one or more direct, and one or more indirect stimulation-induced modulation effects. In the case of indirect modulation effects with directional, rather than binary, outcomes, a) increases in inter-discharge interval and b) decreases in mean electrographic seizure pattern duration, were scored as presence of an effect. We calculated the probability of achieving responder status given the presence of direct modulation effects, and of indirect modulation effects, using two Fisher exact tests, as well as the probability of improving either of the 3 seizure manifestation variables.

## Results

### Identification of electrographic seizure patterns

We analyzed ECoG data from 11 patients implanted with the RNS System (3 male, mean age = 35 years (range 19-65), average duration of epilepsy = 19 years (range 5-37)). The mean time after transplantation to activation of responsive stimulation was 22 weeks (SD: 8.26). One patient was excluded because stimulation was never turned on, as post-surgically he spontaneously became seizure free. Patient 3 underwent resection and electrode repositioning 12 months post-implantation, and post-resection data were excluded. Diagnostic evaluation revealed MRI-appreciable structural abnormalities in 5 patients (Figure 1, Supplementary Figure 1, Table 1). A total of 14,634 ECoG files were visually reviewed, corresponding to 170 months of post-surgical implantation recordings spanning a 34-month total study period, and a total of 5,148 seizure electrographic patterns were identified. We marked the onset of electrographic seizure patterns (Figure 1b, c, e, f, top) and created aligned 3D time-frequency power plots for each event (Figure 1b, c, e, f, middle), which were in turn averaged to reduce individual sample variations and random incidental noise (Figure 1b, c, e, f, bottom).

The spectral features of electrographic seizure patterns per programming epoch, during which stimulation parameters remained constant, were identified. For every patient, and per programming epoch, we visually evaluated all detected events in both time and frequency spectrum domains. We discovered two main categories of neuronal electrical stimulation effects: 1) direct effects, where the recorded events manifested systematic time and/or frequency changes in the immediate post-stimulation interval, and 2) indirect effects, where the recorded events manifested time and/or frequency changes before and/or long after stimulation, and/or could not be attributed to direct stimulation.

### Stimulation can inhibit electrographic seizure patterns

We first characterized the modulation of electrographic seizure patterns that resulted from individual stimulation events. In four patients, we observed electrographic seizure patterns whose progression was terminated in the immediate post-stimulation period, shortly after (<5 s) the application of the first stimulation pulse, following which the ECoG returned to the interictal background level (Figure 2a). This effect, which we termed direct inhibition, is consistent with that expected from prior literature (Kossof *et al*., 2004). In two of these patients, we also identified a variation of this effect, where activity subsequently rebounded and evolved into a typical electrographic seizure pattern despite transient suppression (Figure 2b). Direct electrographic seizure pattern inhibition emerged at a mean of 14 weeks (SD: 11.72) after stimulation was activated.

**Figure 2.**
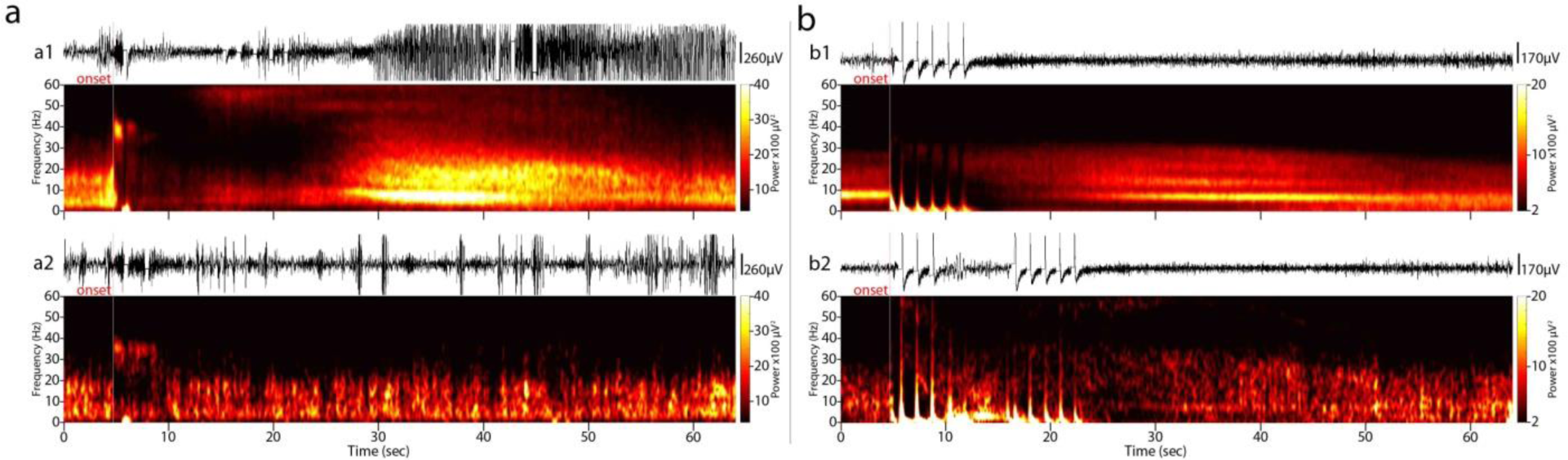
Direct inhibition of electrographic seizure patterns - the current hypothesis for the RNS principle of operation. (a) Time-paired raw ECoG and averaged spectrogram of stimulated electrographic seizure patterns for patient 3/non-responder; most electrographic seizure patterns evolved despite stimulation (a1, average spectrogram of n=62 electrographic seizure patterns; channel 4; weeks 9-20), and some were inhibited shortly after their onset (a2, n=9; channel 4; weeks 9-20), the latter in both cases marked by an abrupt suppression of the normal background band and the appearance of a distinctive high beta 30-40Hz oscillation (red and white lines over the time and frequency domains, respectively). (b) A variation of direct inhibition was observed in patient 2/non-responder, where among the regular electrographic seizure patterns (b1, n=118; channel 2; weeks 26-46) we identified some that were temporarily inhibited by stimulation before resuming again and developing into a typical electrographic seizure pattern (b2, n=6; channel 2; weeks 26-46). Direct electrographic seizure pattern inhibitions emerged at a mean of 14 weeks (SD: 11.72) after stimulation was activated.

### Stimulation can transiently modulate the frequency content of electrographic seizure patterns

We also identified changes in the frequency content of electrographic seizure patterns that resulted from individual stimulation events. In three patients, we observed modulation of the spectral constituents of electrographic seizure patterns that occurred shortly after (mean <5 s) the application of the first stimulation pulse and/or during the stimulation interval (Figures 3 and 4b2, Supplementary Figure 3). This direct frequency modulation effect was variable in nature and consisted of both attenuation of the baseline frequencies (Figure 3a-f, Supplementary Figure 3a-c, right), as well as the genesis of novel oscillations at higher-than-baseline frequencies (Figure 3b-g, right). Note in the case of patient 2, the newly appearing gamma-range (55-60Hz) oscillation persisted in time both during the development of the electrographic seizure patterns per programming epoch and across programming epochs (Figure 3b-g, left). In patient 5, with typical electrographic seizure patterns in the theta-alpha range (5-13Hz, Figure 4b1), we observed post-stimulus frequency modulation into a wide-band diffuse pattern (4-55Hz, Fig. 4b2), first observed during the 5th week of stimulation. Except for patient 5’s electrographic seizure patterns, the baseline onset frequency reappeared after several programming epochs (Figure 3g, Supplementary Figure 3d-e). In patient 2, the frequency modulation was additionally observed in electrographic seizure patterns that did not receive triggered stimulation, suggesting the presence of underlying chronic stimulation effect (Figure 3h). Direct frequency modulation effects emerged at an average of 12.1 weeks (SD: 10.5) following activation of responsive stimulation.

**Figure 3.**
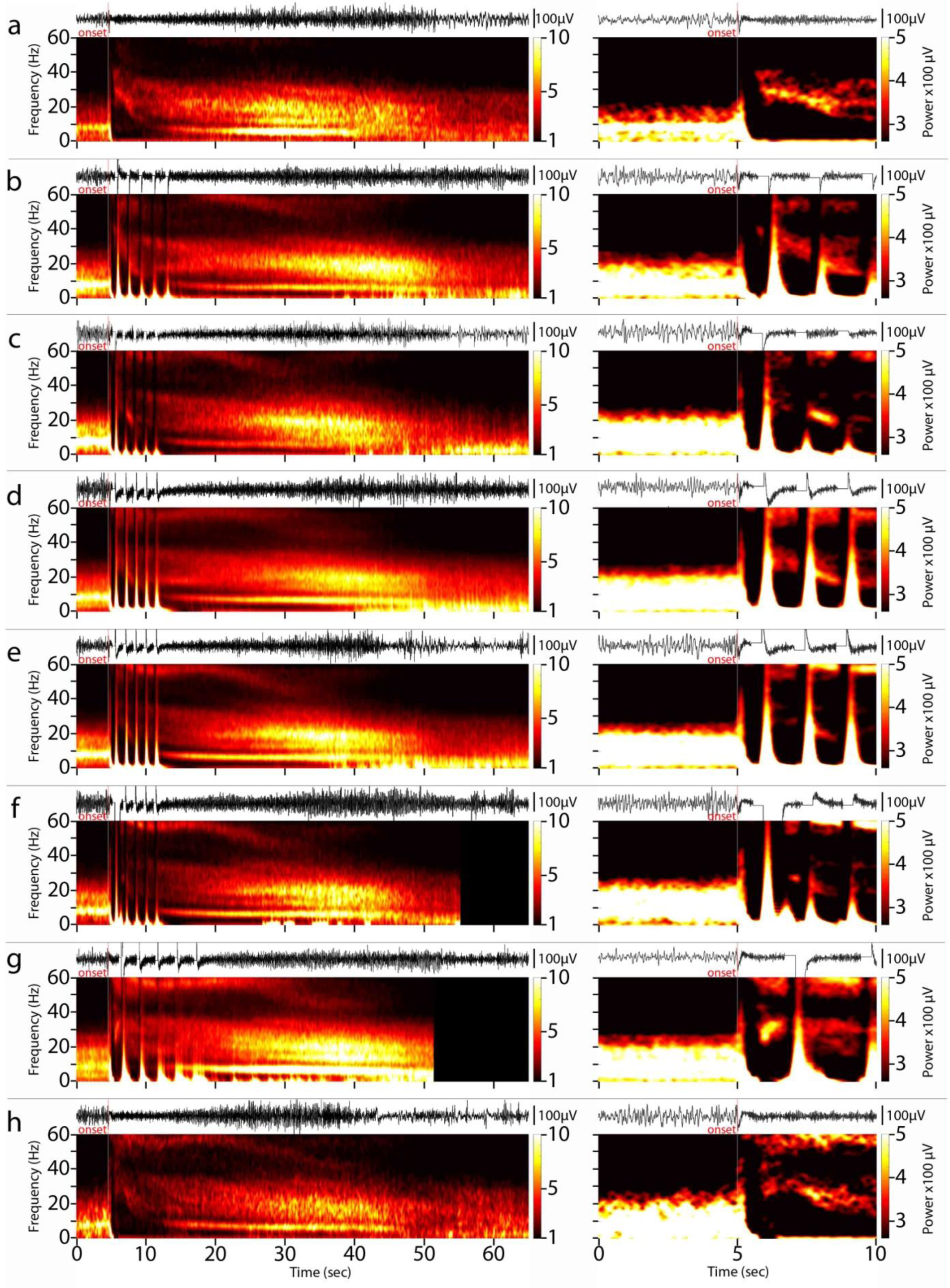
Direct frequency modulation. (a) Time-paired raw ECoG and averaged spectrum plots of typical baseline non-stimulated electrographic seizure patterns (left) of patient 2/non-reponder, zoomed in time and power-wise enhanced symmetrically around the onset (right). Notice the high-to-low beta (from 40 to 20Hz) frequency band systematically featuring at the electrographic seizure patterns’s onset across samples (n=24; channel 2; baseline weeks 0-5). (b-g) Respective figures for the consecutive programming epochs (channel 2; weeks 5-16: n=46, weeks 16-26: n=34, weeks 26-46: n=125, weeks 46-64: n=76, weeks 64-73: n=27, weeks 73-97: n=66; the full 60 s post onset interval was not available for the last 2 programming epochs due to limitations of the device). Notice the progressive peri-stimulus attenuation of the initially dominant beta oscillation (b to f, right), and the equally progressive genesis of a novel gamma (55-60Hz) oscillation in its stead. The newly appearing gamma-range oscillation persisted in time both during the development of the electrographic seizure patterns per programming epoch and across programming epochs. (h) Both frequency bands appear concurrently during the last programming epoch (n=73; channel 2; weeks 73-97), as well as in electrographic seizure patterns that were missed by stimulation throughout the stimulation periods (n=12; channel 2; weeks 5-97). The latter observation suggests the presence of underlying chronic stimulation effects.

**Figure 4.**
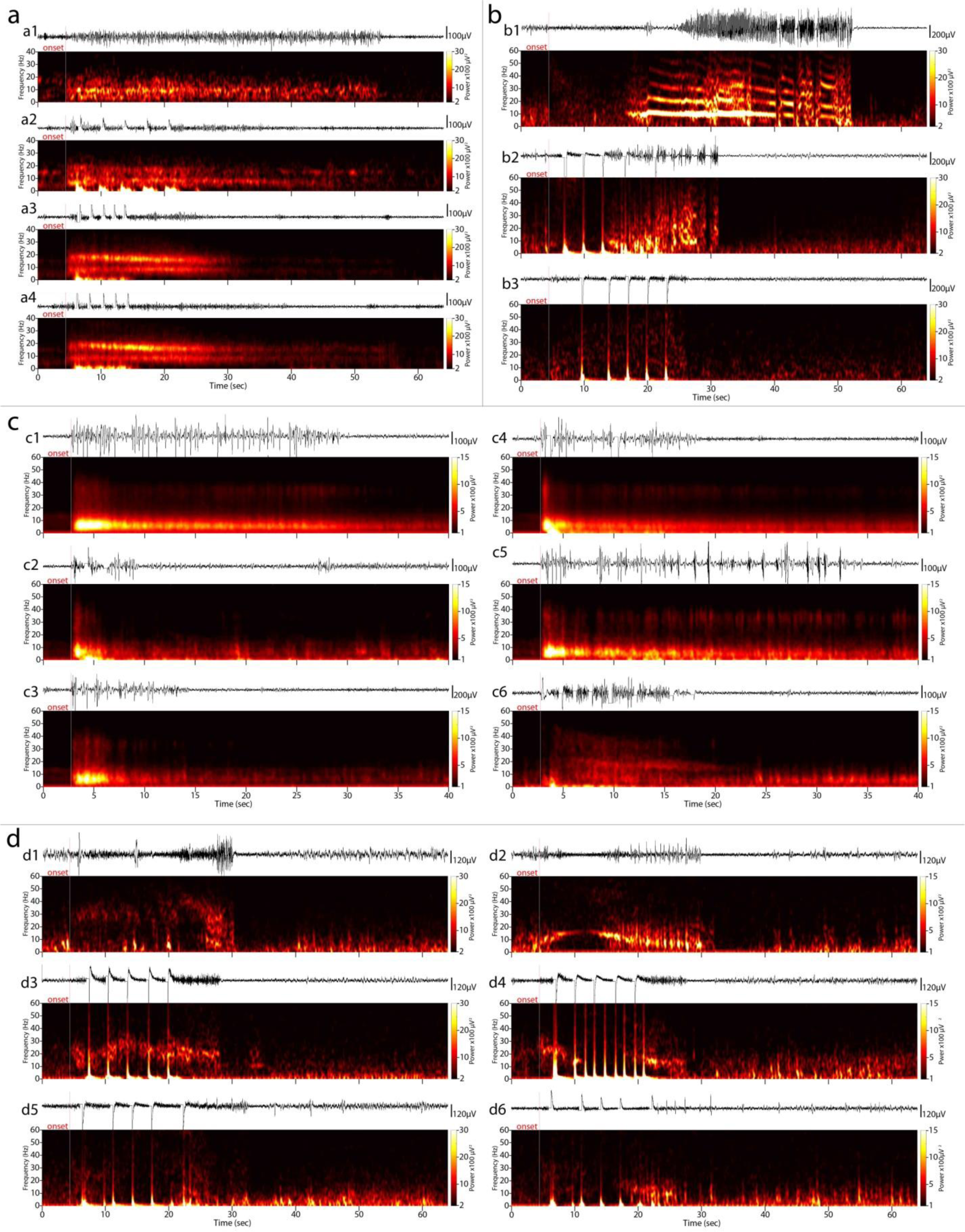
Indirect frequency modulation of electrographic seizure patterns. (a) Patient 7/responder: Subsequent pairs of raw ECoG and averaged spectrograms of electrographic seizure patterns from the baseline weeks 0-5 (a1), weeks 5-11 (a2), weeks 19-24 (a3), and weeks 62-71 (a4) (channel 2; n=2, 2, 39, and 34, respectively). Notice the baseline alpha-range (9-10Hz) rhythm being replaced by a double-band of independent theta and beta frequencies (6-10 Hz and 13-20 Hz respectively), with the latter exhibiting a progressive tendency to dominate the electrographic seizure patterns. The patient also had a mean 66% reduction in total electrographic seizure pattern duration since stimulation week 12 (a3). (b) Patient 5/responder: Subsequent pairs of raw ECoG and averaged spectrograms of electrographic seizure patterns from the baseline weeks 0-7 (b1), featuring continuous oscillations in the theta-alpha range (5-13Hz), and weeks 7-22 (b2, b3) (channel 4; n=2, 3, and 3, respectively). Notice the two clusters of electrographic seizure patterns during the first stimulation period, featuring diffuse spectral content (4-55Hz), from which the baseline dominant ictal alpha rhythm is absent. Although the first cluster (b2) shows frequency modulation after the application of the first pulse and during the stimulation interval, therefore categorized as a direct frequency modulation effect, the second cluster (b3) does not show direct changes but rather a pre-stimulation established continuously diffuse high-frequency pattern. The patient also had a mean 43% reduction in total electrographic seizure patterns duration since the 3rd week of stimulation. (c) Patient 8/responder: Subsequent pairs of raw ECoG and averaged spectrograms of electrographic seizure patterns from the baseline weeks 0-10 (c1), with baseline electrographic seizure patterns characterized by a semi-continuous progression of discharges in the delta-to-alpha range (2-10 Hz), weeks 10-19 (c2, c3), and weeks 28-43 (c4, c5, c6) (channel 2; n=186, 46, 51, 253, 51 and 22, respectively). Notice, 1: the initially strong direct inhibitory stimulation effect, with onset during the 4th week of stimulation, (c2) being progressively ineffective (c4); 2: the initially spontaneous reduction in electrographic seizure patterns duration (c3) being progressively obsolete (c5), as highlighted by electrographic seizure patterns that missed stimulation; and 3: a cluster of electrographic seizure patterns where the characteristic ictal theta discharges are absent, and feature a wide-band high-frequency (4-50Hz) content instead. The latter effect was first observed during the 20th week of responsive stimulation. (d) Patient 1/responder, exhibited different electrographic seizure patterns per RNS lead as they were implanted in different parts of the lesion (Supplementary Figure 1): Subsequent pairs of raw ECoG and averaged spectrograms of electrographic seizure patterns from the baseline weeks 0-1 (d1, d2), weeks 1-16 (d3, d4), and weeks 16-24 (d5, d6) (channels 2 and 3; n=4, 5, and 6, respectively). One baseline pattern was that of high frequency (22-53 Hz, d1), and the other was characterized by an arc-like pattern in the delta to low beta range (2-18 Hz, d2). Over subsequent stimulation periods (1-24 weeks), these patterns underwent progressive changes in their spectral content (d3-d6), evolving to diffuse spectrum low power discharges (5-60 Hz and 7-38 Hz, respectively). Overall, indirect frequency modulation effects emerged at an average of 21.7 weeks (SD: 25.9) following activation of responsive stimulation.

### Spontaneous attenuation of electrographic seizure patterns

Next, we characterized indirect effects, where recorded events manifested evidence of modulation that could not be attributed to a specific stimulation event. In one patient, we observed seizure patterns whose progression was spontaneously discontinued in the absence of a direct stimulation event, during periods of baseline activity, i.e. long after (>27sec) the application of the nearest initial stimulation pulse and end of subsequent triggered stimulation pulses (>11sec) (Figure 5). We termed this effect indirect attenuation. The typical electrographic seizure patterns began with a diffuse electro-decrement followed by the development of an ictal theta-range (4-8 Hz) rhythm (Figure 5a1-d1) evolving into high amplitude/power paroxysmal wide-band delta-to-beta-range (2-30 Hz) rhythms overlaid with higher gamma (>30Hz) frequencies. Spontaneous discontinuation of these electrographic seizure patterns (Figure 5c1, c2) occurred at the point of transition from the singular theta to the wide-band ictal pattern. The onset of this spontaneous indirect effect was first observed during the 9th week of stimulation and reappeared regularly up to time of last follow-up (112th stimulation week).

**Figure 5.**
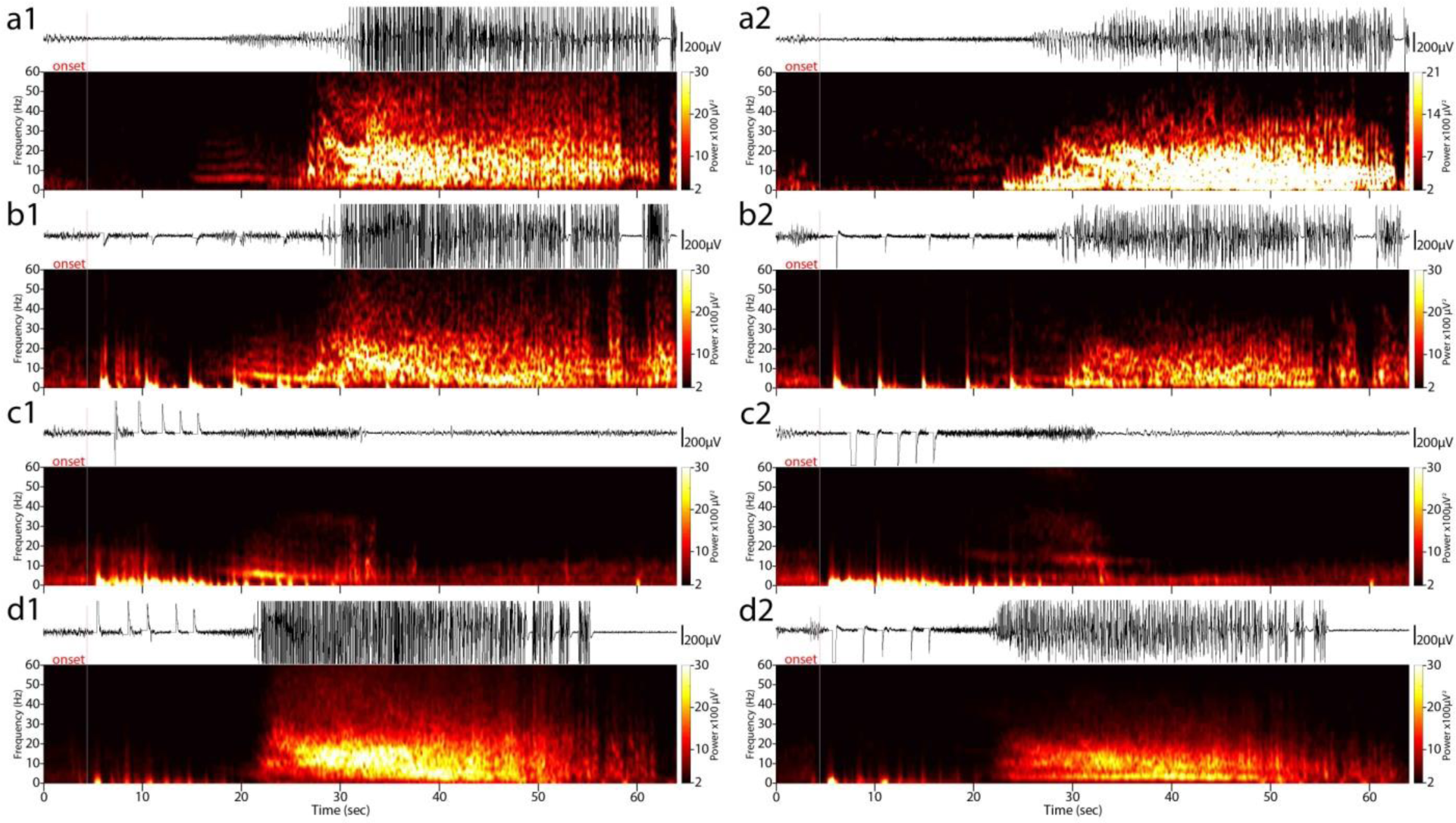
Indirect spontaneous attenuation of electrographic seizure patterns. (a1) Time-paired raw RNS data and averaged time-frequency power plots of electrographic seizure patterns from the free of stimulation baseline phase (n=2; channel 1; weeks 0-1) for patient 11/responder. (a2) shows channel 2 from the same period. These electrographic seizure patterns start with a diffuse electro-decrement, that is, a reduction in amplitude/power in all frequency bands, followed by the development of a theta-range (4-8 Hz) rhythm (most apparent on channel 1) evolving into high amplitude/power paroxysmal wide-band delta-to-beta-range (2-30 Hz) rhythms overlaid with higher gamma (>30Hz) frequencies. (b1, b2) Channels 1 and 2, respectively, during weeks 3-7, showing the same onset pattern occurring under stimulation conditions (n=4). (c1, c2) From weeks 7 to 112, a small but clearly distinct number of electrographic seizure patterns were observed, in which the development of the ongoing activity was spontaneously interrupted before the genesis of the delta waves (n=20), and the ECoG returned to normal background levels; note that the breakdown occurs more than 27 s after the 1st stimulation pulse and the bulk of stimulation ends almost 11 s before this spontaneous inhibition. Spontaneous discontinuation of these electrographic seizure patterns occurred at the point of transition from the singular theta to the wide-band ictal pattern. The onset of this spontaneous indirect effect was first observed during the 9th week of stimulation and reappeared regularly up to time of last follow-up (112th stimulation week). (d1, d2) Electrographic seizure patterns from weeks 42-65, where their mean duration is significantly reduced (37% reduction); the leading theta rhythm following the diffuse electro-decrement onset is less apparent among the co-existing paroxysmal alpha and higher frequencies (n=14).

### Stimulation can induce persistent frequency modulation of electrographic seizure patterns

We also identified changes in the spectral constituents of electrographic seizure patterns that were not related to individual stimulation events, i.e. indirect frequency modulation, which emerged over time during the course of RNS treatment in four patients. The narrow alpha-band (9-10Hz) electrographic seizure patterns of patient 7 (Figure 4a1) transformed into a double-band pattern of concurrently evolving theta and beta frequencies (6-10 Hz and 13-20 Hz respectively, Figure 4a2-4). Patient 5, with similar typical electrographic seizure patterns containing continuous oscillations in the theta-alpha range (5-13Hz, Figure 4b1), exhibited an indirect diffuse spectral (4-60Hz) low amplitude/power pattern that was present prior to stimulation pulses (Figure 4b3), first observed during the 9th week of responsive stimulation. Patient 8, with baseline electrographic seizure patterns characterized by a semi-continuous progression of discharges in the delta-to-alpha range (2-10 Hz, Figure 4c1), and an onset of directly inhibited electrographic seizure patterns during the 4th week of stimulation (Figure 4c2), developed completely distinct wide-band (4-50 Hz) electrographic seizure patterns that were independent of specific stimulation pulses. This effect was first observed during the 20th week of responsive stimulation. Patient 1, exhibited baseline electrographic seizure patterns that were different between the two RNS leads (Figure 1, Supplementary Table 1). One pattern was one of high frequency content (22-53 Hz, Figure 4d1), and the other was characterized by an arc-like pattern in the delta to low beta range (2-18 Hz, Figure 4d2). Over subsequent stimulation periods (1-24 weeks), these patterns underwent progressive changes in their spectral content (Figure 4d3, d4), evolving to diffuse spectrum low power discharges (5-60 Hz and 7-38 Hz respectively, Figure 4d5, d6). These frequency modulation effects appeared during the first week of responsive stimulation.

### Stimulation can increase the refractory interval between ictal spikes

In one of our patients we observed indirect fine fragmentation of electrographic seizure patterns, in which the refractory interval between consecutive seizure spike discharges increased (baseline interval mean ≈ 200ms, post-effect interval mean ≈ 800 ms) (Figure 6), independent of the timing of stimulation onset. Although increased inter-discharge intervals were observed sporadically during baseline and previous programming epochs (Figure 6a, b), they were only established as the dominant electrographic seizure pattern after the 60th week of stimulation (Figure 6c, d).

**Figure 6.**
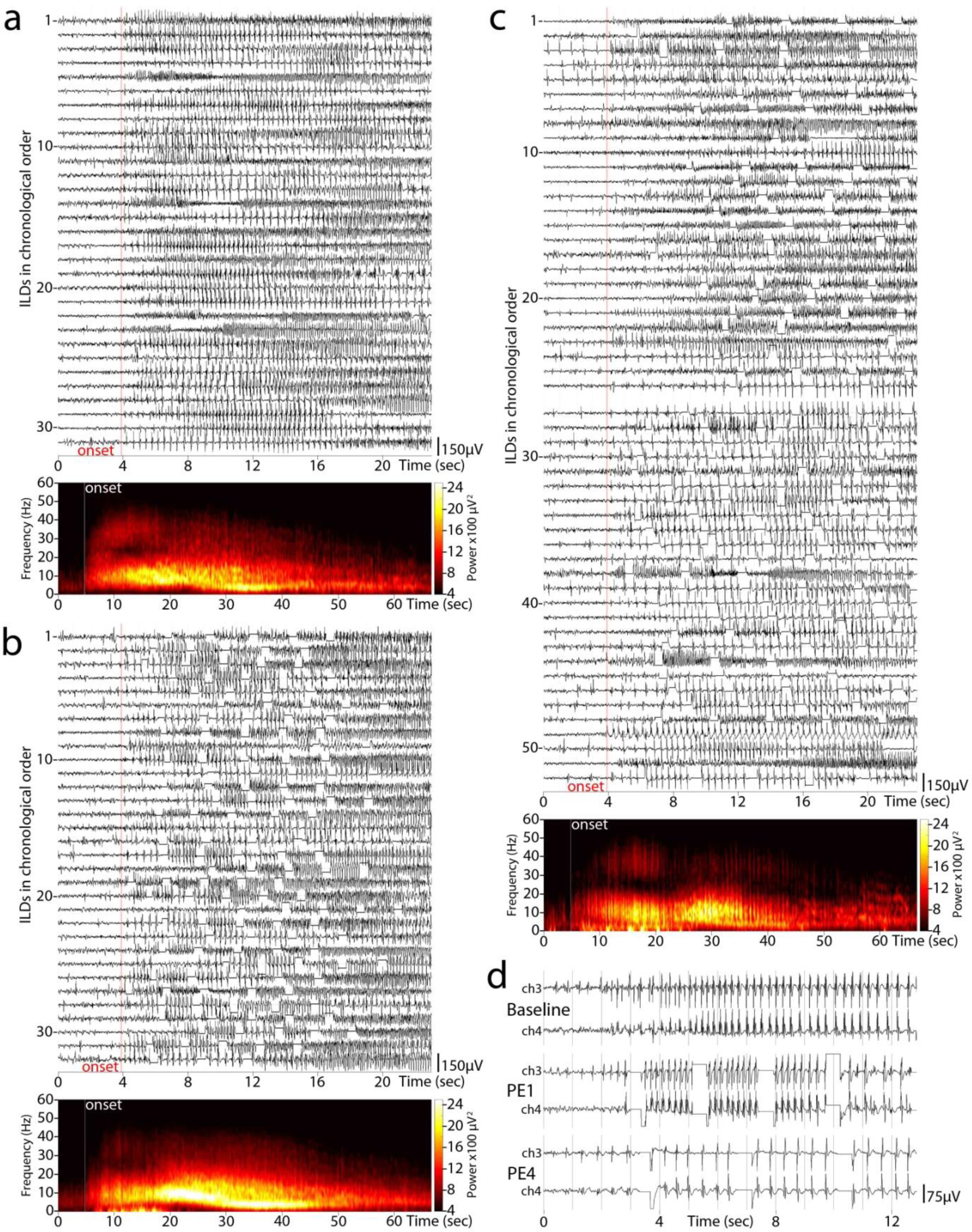
Indirect fine fragmentation of electrographic seizure patterns - inter-discharge interval modulation. (a) Baseline weeks 0-5 electrographic seizure patterns recorded on channel 3 of patient 6, appearing in chronological order from top to bottom (n=32). An average time-frequency plot, not time-paired to the time domain above, is shown below. (b) Programming epoch weeks 5-16 electrographic seizure patterns, also in chronological order (n=32), accompanied by a not time-paired averaged spectrogram. No notable differences are observed between the two periods, neither in the time nor in the frequency domain. Same patterns appeared during weeks 16-34 and 34-43 (not shown). (c) Programming epoch weeks 43-66 electrographic seizure patterns (n=52): After the first half of this programming epoch, electrographic seizure patterns appeared with significant increase in inter-discharge interval (from a previous baseline mean of ≈200 ms, to a post-effect interval of ≈800 ms); the averaged spectrogram below corresponds to electrographic seizure patterns of the programming epoch’s second half (n=27), assuming a comb-like pattern due to the low amplitude inter-discharge intervals introduced. (d) Note that increased inter-discharge interval discharges appear sporadically and briefly during the previous programming epochs but were only established as the dominant electrographic seizure pattern after the 60th week of stimulation.

### Stimulated electrographic seizure patterns can become fragmented

In two patients, we observed examples of indirect coarse fragmentation of electrographic seizure patterns, in which the continuity of an ongoing discharge was interrupted by segments of normal background activity (Figure 7, Supplementary Figure 4). These seizure fragments occurred in random intervals from the onset of any given stimulation and with variable duration. Both patients’ baseline electrographic seizure patterns were comprised of low frequency theta/delta content with spikes. In patient 10, although directly inhibited electrographic seizure patterns appeared as early as the 5th week of stimulation and where present throughout the subsequent programming epochs (up to 105 weeks, Figure 7i), fragmented electrographic seizure patterns first appeared during the 8th week of stimulation and persisted equally throughout (Figure 7c-g). In patient 8, fragmented electrographic seizure patterns first appeared during the 19th week of stimulation and persisted up to the 5th week of stimulation. The mean onset of the fragmentation effect was 13 weeks (SD: 7.77) after stimulation activation.

**Figure 7.**
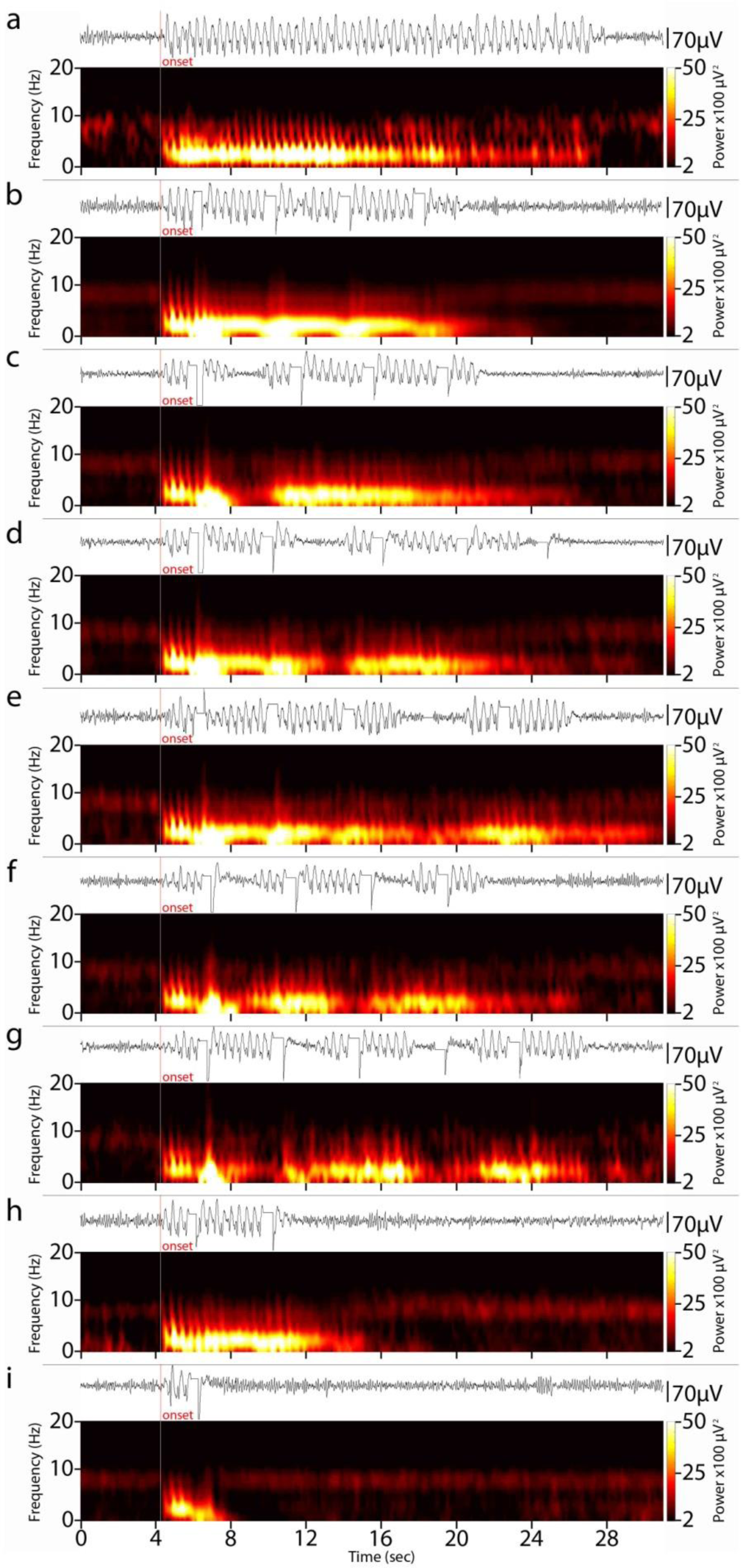
Indirect coarse fragmentation of electrographic seizure patterns. (a) Baseline weeks 0-7 raw ECoG and time-paired averaged spectrogram (n=3; channel 1) showing the typical time and frequency electrographic seizure patterns of patient 10, comprised of a basic low theta/high delta spike-wave discharge sequence. (b) Stimulated electrographic seizure patterns from programming epoch weeks 13-19 (n=100). (c-g) Beginning from programming epoch weeks 24- 30, RNS recorded electrographic seizure patterns with pronounced discontinuities during their evolution, during which the ECoG returned to the background levels, thereby resulting in a variably fragmented pattern. For the purpose of demonstrating both the fragmentation effect as well as its variable character, we arbitrarily clustered electrographic seizure patterns into five categories (channel 1; weeks 54-90): (c) patterns with a single fragmentation between 0-5 s after the onset (n=41); (d) patterns with a single fragmentation between 5-10 s after onset (n=36); (e) patterns with a single fragmentation more than 10 s after onset (n=20); (f) patterns with double fragmentation between 0-5 s and 5- 10 s after onset (n=16); (g) patterns with double fragmentation between 0-5 s and more than 10 s after onset (n=8). (h) The patient manifested a subgroup of electrographic seizure patterns with a mean 55% reduction in duration (n=24; channel 1; weeks 54-90). (i) Directly inhibited electrographic seizure patterns were also present (n=47; channel 1; weeks 54-90). Directly inhibited electrographic seizure patterns appeared as early as the 5th week of stimulation, while fragmented and shortened electrographic seizure patterns first appeared during the 8th and 53rd week of stimulation, respectively.

### Stimulation modulates the mean electrographic seizure pattern duration

Finally, we observed significant bi-directional changes in the mean duration of electrographic seizure patterns that occurred in the absence of direct stimulation events, in five patients. Patient 5 had a mean 43% reduction in total electrographic seizure patterns duration during weeks 9 to 22 (Figure 4b), patient 7 had a mean 66% reduction in total electrographic seizure pattern duration during stimulation weeks 12 to 86 (Figure 4a3), patient 10 manifested a subgroup of electrographic seizure patterns with a mean 55% reduction in duration during stimulation weeks 53 to 105 (Figure 7h), and patient 11 had a mean 37% reduction in electrographic seizure patterns duration during weeks 81 to 112 (Figure 5d1, d2). In contrast, patient 9 progressively reached a mean 132% increase in electrographic seizure pattern duration by stimulation week 7 (Supplementary Figure 5). Although stimulation-induced direct inhibition reduces the duration of the ictal pattern (Figure 7b, h, and Figure 4c4, c5), the indirect reduction in electrographic seizure pattern duration is a different phenomenon as the offset of the electrographic seizure patterns is not related to the offset of any of the applied stimuli. The mean onset of effects that altered ictal duration was 31 weeks.

### Indirect modulation effects correlate with improved clinical outcome

To determine whether direct and/or indirect modulation effects were correlated with the reduction of seizures, Fisher exact tests were performed to separately measure the probability of observing either type of effect and achieving responder status (defined as an outcome of Engel class III or better, Table 1). The odds ratio (OR) for indirect modulation effects was infinity (p=0.02), demonstrating a strong and significant association between the presence of one or more indirect modulation effects and good clinical response. The OR for direct effects was not significant (0.67, p=1.0). Fisher exact tests then were performed to measure the probability of observing a direct or indirect modulation effect and achieving reduction in either seizure frequency, severity, or duration. The OR for indirect modulation effects was significant for each of these outcomes (seizure occurrence frequency: OR=infinity, p=0.0047; seizure severity: OR=infinity, p=0.007; seizure duration: OR=28, p=0.032) while the OR for direct modulation effects was not significant for any outcome (seizure frequency: OR=0.67, p=1.0; seizure severity: OR=0.0, p=0.1; seizure duration: OR=0.25, p=0.56). The mean onset of modulation effects was 11.5 stimulation weeks (SD: 3.5) for direct effects and 24 weeks (SD: 22.24) for indirect effects, suggesting that network plasticity is required for both types of changes.

## Discussion

We investigated the validity of the direct inhibition hypothesis in 11 patients implanted with RNS. We extracted their data from the PDMS and visually reviewed all files to mark the onset of electrographic seizure patterns. In turn, we clustered electrographic seizure patterns per programming epoch and generated averaged 3D time-frequency power graphs. Combining the visually appreciated features in the time and frequency domains, we conducted a novel evaluation of the electrophysiological effects of closed-loop stimulation. We discovered two categories of stimulus-related modulation effects: 1) direct inhibition of electrographic seizure patterns, in accordance with traditional hypotheses of RNS mechanism of action, and 2) direct frequency modulation; a novel finding of direct post-stimulus changes in the spectral content/signature of the electrographic seizure patterns that includes attenuation of prominent baseline bands (that may rebound in time) as well as the genesis of completely new electrographic seizure patterns. These direct stimulation effects did not correlate with clinical outcomes. The most important finding in this study is the discovery of indirect modulation effects, which occurred independent of specific stimulation events and which were strongly correlated to clinical outcomes.

Evaluation of time-frequency features beyond the narrow direct stimulation time interval, throughout the electrographic seizure patterns’ time-course and across programming epochs, revealed five categories of indirect modulation effects: 1) spontaneous inhibition, where electrographic seizure patterns were interrupted long after the applied stimulation pulses, 2) frequency modulation, where the spectral signature/pattern showed remarkable changes in frequency content, 3) fine fragmentation, where the refractory period between consecutive ictal spikes was markedly increased, 4) coarse fragmentation, where the discharge continuity was intermittently spontaneously interrupted by brief background intervals that were not a result of direct stimulation, and 5) modulation of the electrographic seizure pattern duration, where the mean interval between the onset and the offset of the electrographic seizure patterns underwent remarkable changes (both reductions and increases) that cannot be attributed to direct inhibition of the electrographic seizure patterns.

The direct effects of electrical stimulation during on-going epileptic activity have been widely described. Penfield was the first to observe and report stimulation-induced inhibitory effects of electrical stimulation on active epileptogenic neural tissue (Penfield and Jasper, 1954), later also verified by stimulation of hippocampal slices in-vitro (Jefferys, 1981; Durand, 1986). Electrical stimulation has also been reported to have a significant inhibitory effect on interictal and ictal cortical activity in epilepsy patients (Velasco *et al*., 2000; Yamamoto *et al*., 2002; Kinoshita *et al*., 2004; Kossoff *et al*., 2005; Kinoshita *et al*., 2005; Yamamoto *et al*, 2006;Elisevich *et al*., 2006; Velasco *et al*., 2009; Child *et al*., 2014; Ludstrom *et al*., 2016, 2017; Valentin *et al*., 2017; Kerezoudis *et al*., 2017). The RNS system is a valuable surgical option for patients refractory to both AEDs and traditional epilepsy surgery (Stacey and Litt, 2008), with data supporting its superiority to medical management (Heck *et al*., 2014; Geller *et al*., 2017; Jobst *et al*., 2017). However, apart from the few original publications (Lesser *et al*., 1999; Kossoff *et al*., 2004; Osorio *et al*., 2005), little is known about how closed-loop stimulation affects the time-course of the detected electrographic seizure patterns and related mechanisms of action for reducing seizure severity and frequency. The previously assumed mechanism is a direct one, by which the application of an electric pulse close to the origin of the electrographic seizure pattern interrupts its evolution and returns the ECoG background to its interictal state (Kossoff *et al*., 2004; Skarpaas and Morrell, 2009; Morrell and Halpern, 2016). The main hypothesis behind direct inhibition of on-going epileptic activity is that the electric stimulation current transiently activates local post- synaptic potentials (PSPs) that create extracellular fields opposing and partially or fully cancelling those created by the epileptogenic excitatory post-synaptic potentials (EPSPs). In this model, as extracellular fields are essential for synchronization between distant neuronal pools, stimulation reduces the ability of the underlying tissue to synchronize its excitatory neuronal population activity, thereby acting as a neuronal de-synchronizer (Durand, 1986).

Our indirect modulation findings, whose manifestation cannot be adequately explained in direct response terms, expand an alternate hypothesis. All indirect modulation patterns did not appear during the stimulation period and did not alter their main features after completion of stimulation. Indeed, the indirect frequency modulation effects, as well as the direct variety, show that parts of the underlying epileptogenic tissue can be “forced” by stimulation to oscillate at different frequencies than before. One explanation for these findings is that stimulation establishes extracellular electric field barriers between functionally interconnected epileptogenic populations, thereby isolating excitatory neuronal pools. These neuronal pools become separated and independent from the core epileptogenic pool over time resulting in 1) lower amplitude/power of baseline oscillations (progressive attenuation) due to the reduced number of participating neuronal pools, and 2) the appearance of higher frequency oscillations (representing EPSP activity) due to multiple spatially scattered populations unable to achieve high levels of synchronization (Bragin *et al*., 2000; Schevon *et al*., 2008). In general, frequency modulation effects indicate that stimulation can drive parts of the underlying epileptogenic network to synchronize at alternative frequencies. On the other hand, a progressive failure of excitatory neuronal populations to achieve sufficiently high levels of synchronization could account for the observed seizure attenuation affect. Likewise, fine and coarse fragmented electrographic seizure patterns can be viewed respectively as manifestations of consecutive and terminal synchronization failures during the development of epileptic excitatory post-synaptic potentials.

Experimental models of epilepsy have shown that the minimal requirement for epileptogenesis is the formulation of an efficiently interconnected and sufficiently large population of excitatory neurons (Jefferys, 1998). In addition, functional studies has shown that progressive increased connectivity of regional epileptogenic networks in focal epilepsy syndromes is positively correlated with increases in the duration and severity of epilepsy, possibly due to the persistent character of locally generated paroxysmal discharges (Englot *et al*., 2016). Thus, electrical stimulation over long periods of time may progressively disrupt the connectivity of the epileptogenic network and reduce the core synchronized population, rendering the clinical manifestation of seizures less severe or subclinical (Zangaladze *et al*., 2008; Singh *et al*., 2015). Such an effect would correspond to responsiveness to RNS (Heck *et al*., 2014; Geller *et al*., 2017; Jobst *et al*., 2017) and is strongly supported by the positive correlation of the indirect modulation effects with the improved post-implantation outcomes of our patients. Outcome studies have reported that a) the number of RNS responders increases with time, and b) seizure reduction percentages also increase over time (Geller *et al*., 2017; Jobst *et al*., 2017),. As the RNS detector settings do not systematically improve over time (Sisterson *et al*., 2018) - a fact that does not favor the direct stimulation effect hypothesis - the documented time-to-response implies an underlying patient-dependent variation in the duration and amount of stimulation required for the establishment of indirect modulation effects. Despite previous reports of effects of acute and subacute chronic stimulation on both ECoG content and seizure control (Velasco *et al*., 2009; Child *et al*., 2014; Ludstrom *et al*., 2016, 2017; Valentin *et al*., 2017), with background normalization over time, and improved seizure control (Velasco *et al*., 2000; Elisevich *et al*., 2006; Kerezoudis *et al*., 2017), our results show that both direct inhibition by stimulation and direct frequency modulation changes observed during the direct stimulation phase have little effect on outcome.

It is important to note that a limitation of our study is the inherent incompleteness of the RNS ECoG data saved in the PDMS. Both the fact that patients are not fully compliant with data uploads, and the fact that the device consistently overwrites its data banks before the upload, render the stored ECoG recordings a fragmented sample of the actual volume of electrographic seizure pattern events. Also, changes in anti-epileptic medication or other life-style variables of our patients were not controlled for in this study.

These results are the first to demonstrate electrophysiological signatures of therapeutic responses to closed-loop brain stimulation for epilepsy. The fact that indirect modulation effects are strongly correlated with improved seizure control, rather than the effects of direct stimulation on triggered seizures, indicates that neural plasticity is required for a therapeutic response, and constitutes a paradigm shift in thinking about neuromodulation for epilepsy Our findings suggest that chronic electrical stimulation progressively disrupts the connectivity of the epileptogenic network and reduces the core synchronized population, thereby rendering the clinical manifestation of seizures less severe. Alternately, stimulation could drive the topographical tightening of connections that accelerate seizure termination (Khambhati *et al*., 2015). Ongoing work seeks to identify the specific stimulation scenarios that produce chronic neuromodulation effects on seizure networks, in order to improve the therapeutic speed and efficacy of closed-loop brain stimulation.

## Acknowledgements

N.D.S and T.A.W. are trainees in the Physician Scientist Training Program (PSTP) at the University of Pittsburgh School of Medicine. The authors thank NeuroPace, Inc. for assistance with data transfer.

## Supplementary Material

**Supplementary Figure 1.**
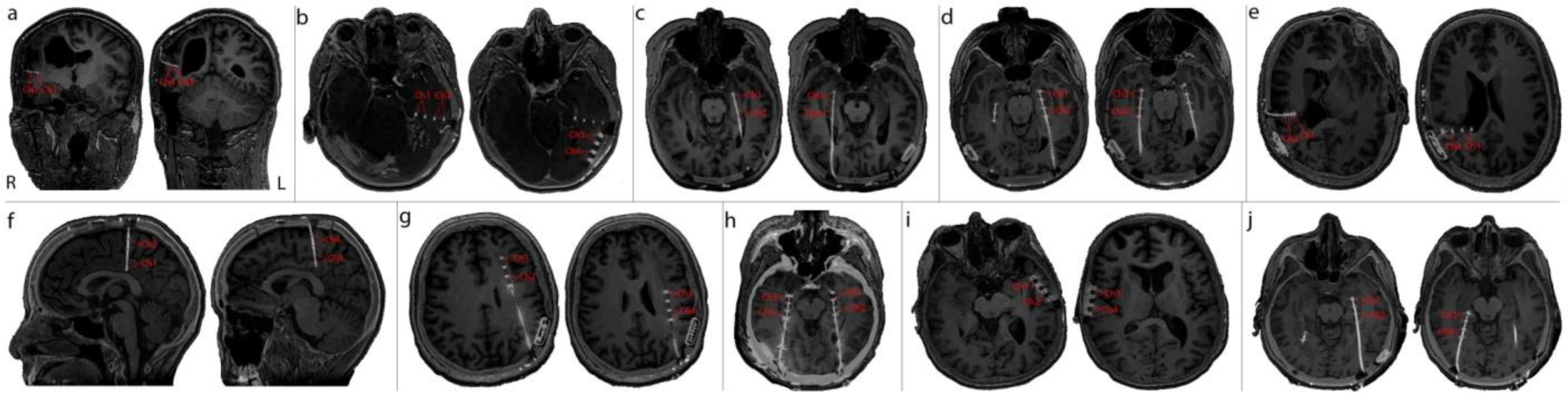
RNS implantation schemes. Fused pre-implantation MR images and post-implantation CT, aligned in over the plane of the trajectory of each implanted RNS lead. (a) Patient 1, with polymicrogyria (PMG) of right posterior fontal/parietal hemispheric distribution. (b) Patient 2, with a left basal temporal arteriovenous malformation (AVM). (c) Patient 3, with bilateral non-lesional mesio-temporal lobe epilepsy (MTLE). (d) Patient 4, with bilateral non-lesional MTLE. (e) Patient 5, with a complex malformation combining PMG and heterotopic components over the posterior frontal/parietal region of the right hemisphere. (f) Patient 7, with non-lesional focal epilepsy involving the mesial structures of the posterior frontal (central) region of the right hemisphere. (g) Patient 8, with an extended heterotopia spanning from the frontal lobe to the parietal area of the left hemisphere. (h) Patient 9, with bilateral non-lesional MTLE. (i) Patient 10, with bilateral non-lesional neocortical temporal lobe epilepsy (TLE). (j) Patient 11, with bilateral heterotopias across the occipito-temporal axis. The implantation scheme of 6 appears in Figure 1. Channels and their derivations appear in red over the fused images for each patient. Orientation follows the radiologic convention.

**Supplementary Figure 2.**
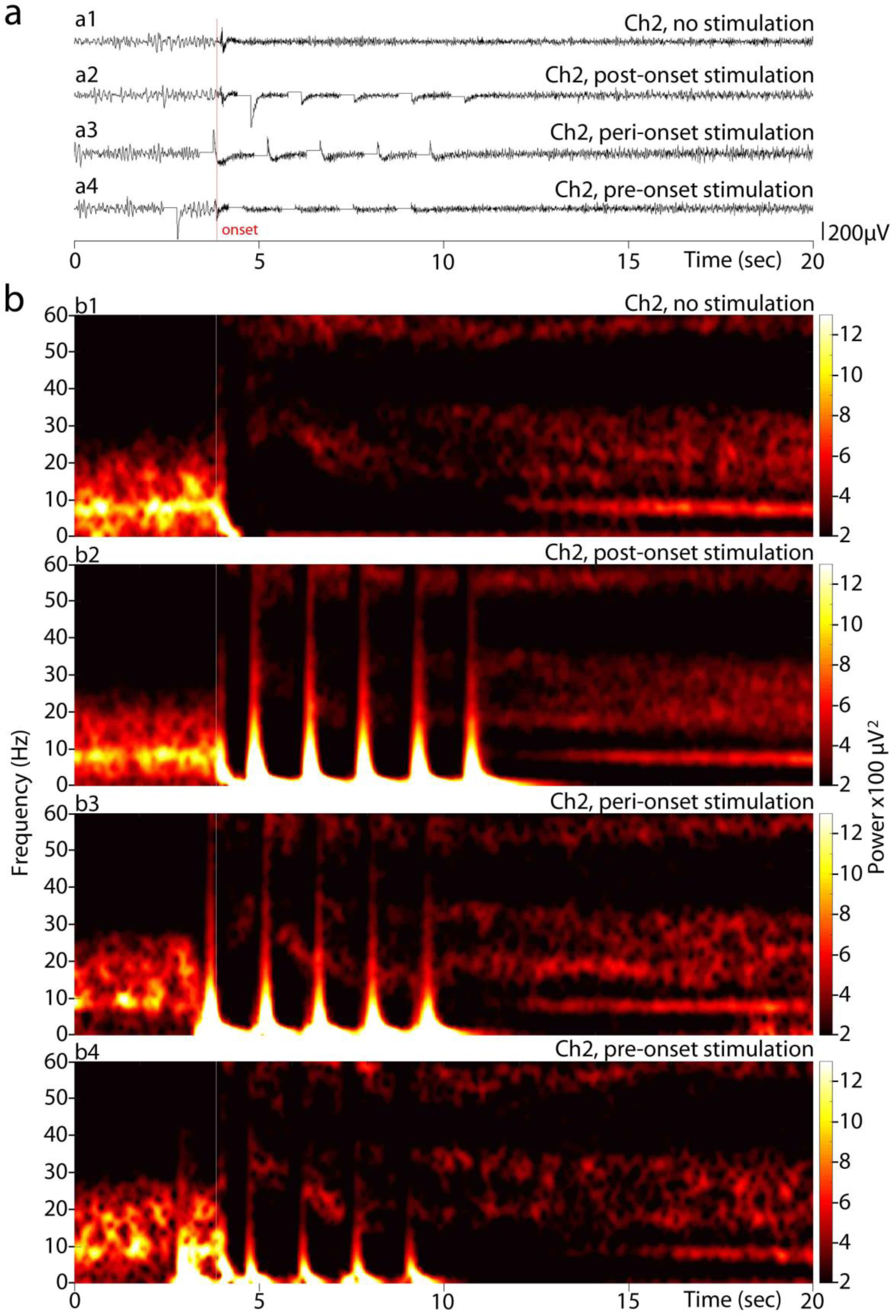
Electrographic seizure patterns relative to the onset of the first RNS stimulation pulse. Visual evaluation of the electrographic seizure patterns across patients resulted in four major clusters, regarding the relative timing between the onset and the application of the first stimulation pulse: (a) Raw RNS-derived ECoGs of patient 2 showing the post-onset, peri-onset and pre-onset stimulated electrographic events (a2-a4), along with a non-stimulated event for reference (a1). (b) Averaged time-frequency power graphs corresponding respectively to the above clusters of the raw ECoGs (n=12, 25, 8, and 5 from b1 to b4 respectively; channel 2; baseline week 0-5 and programming epoch weeks 16-26). Only the former two electrographic seizure pattern categories (non-stimulated and post-onset stimulated events) were included and considered for processing in this study. Electrographic seizure pattern events with peri-stimulus and pre-stimulus onset were not dealt with, in order to maintain both the homogeneity of samples and the accuracy of the spectral averaging processes.

**Supplementary Figure 3.**
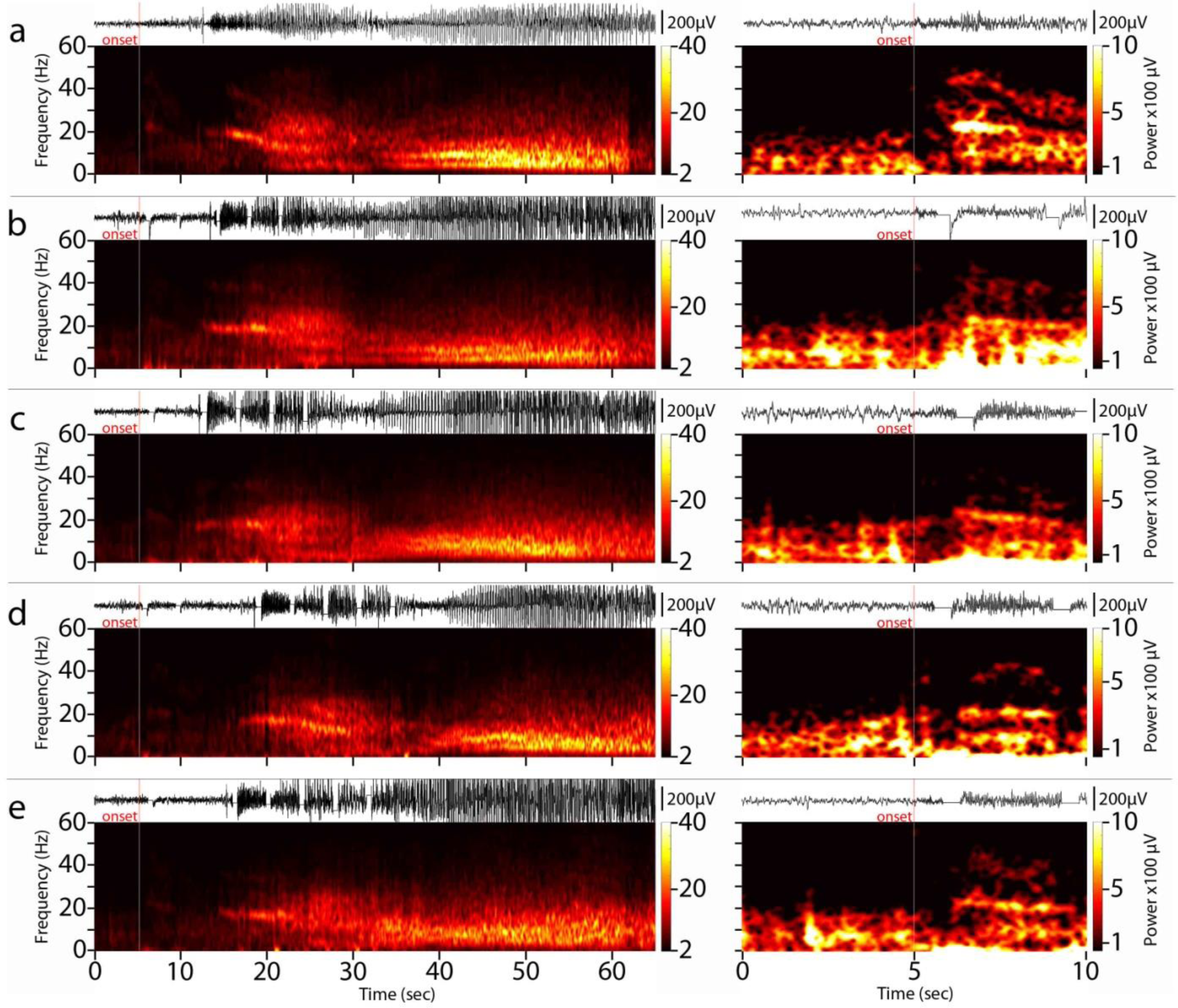
Direct frequency modulation. (a) Time-paired raw ECoG and averaged spectrum plots of typical baseline non-stimulated electrographic seizure patterns (left) of patient 4/non-responder, zoomed in time and power-wise enhanced symmetrically around the onset (right). Notice the beta (from 20 to 35 Hz) frequency band systematically appearing at the onset (n=6; channel 2; baseline weeks 0-8). (b-e), Respective figures for the consecutive programming epochs (channel 1; weeks 8-16: n=12, weeks 16-24: n=14, weeks 24-31: n=14, weeks 31- 45: n=10). Notice the progressive peri-stimulus attenuation of the apparent baseline beta frequency (b to c, right), and its progressive re-appearance in the following programming epochs (d to e, right). Overall, direct frequency modulation effects emerged at an average of 12.1 weeks (SD: 10.5) following activation of responsive stimulation.

**Supplementary Figure 4.**
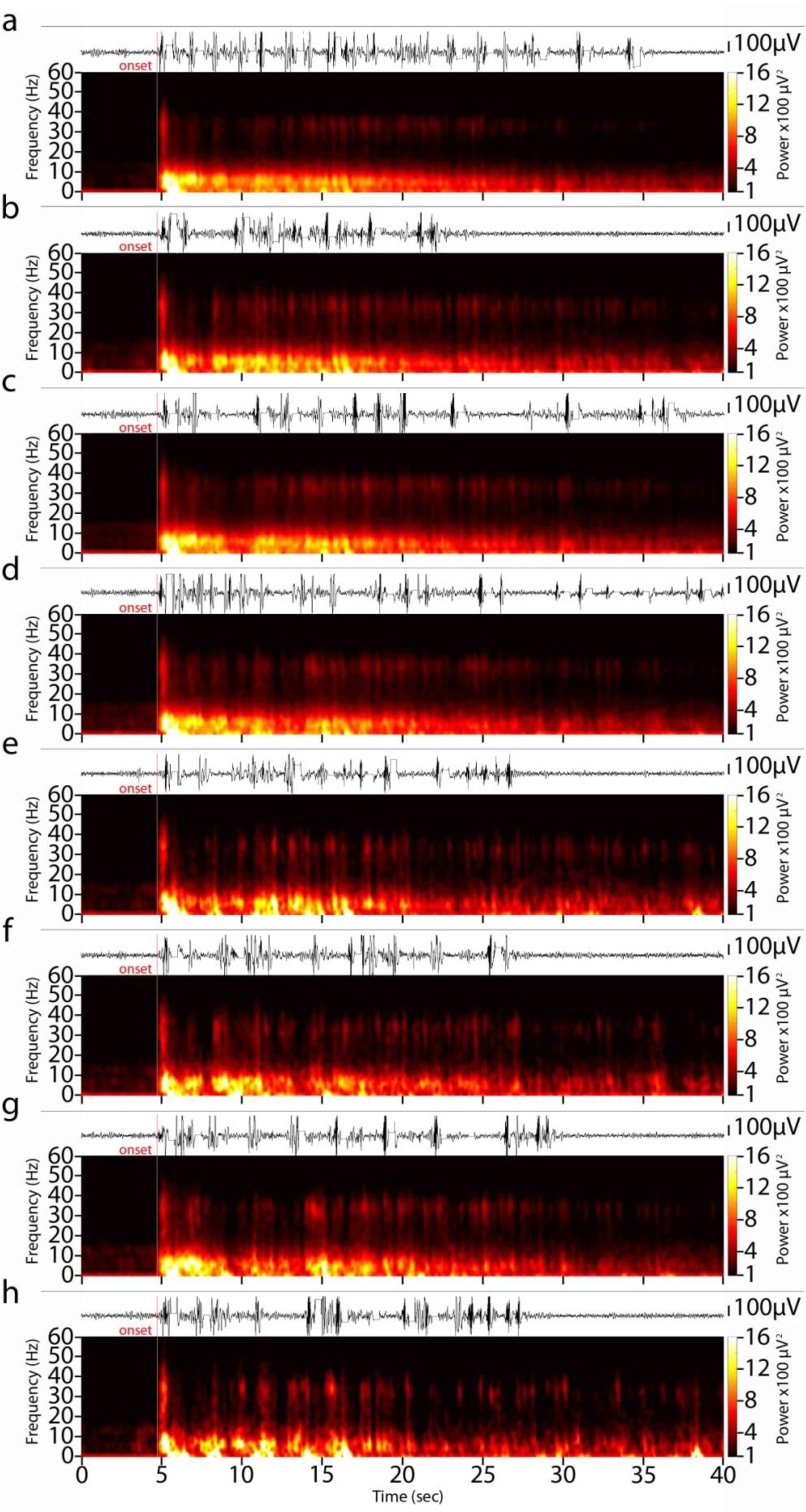
Indirect fragmentation of electrographic seizure patterns. (a) Programming epoch weeks 28- 43 raw ECoG and time-paired averaged spectrogram (n=138; channel 2) showing electrographic seizure patterns of patient 8 that exhibit pronounced discontinuities/fragmentation during their time-course (his baseline electrographic seizure pattern appears in Figure 4c). (b-h) For the purpose of demonstrating both the fragmentation effect as well as its variable character, we arbitrarily clustered these electrographic seizure patterns into seven categories (channel 2; weeks 28-43): (b) patterns with fragmentation between 0-5 s after the onset (n=57); (c) patterns with fragmentation about 5 s after onset (n=88); (d) patterns with fragmentation between 5-10 s after onset (n=66); (e) patterns with double fragmentation between 0-5 s and about 5 s after onset (n=22); (f) patterns with double fragmentation between 0-5 s and 5-10 s after onset (n=19); (g) patterns with double fragmentation between about 5 s and 5-10 s after onset (n=29); (h) patterns with triple fragmentation in all the above arbitrarily selected intervals (n=10). Fragmented electrographic seizure patterns first appeared during the 19th week of stimulation.

**Supplementary Figure 5.**
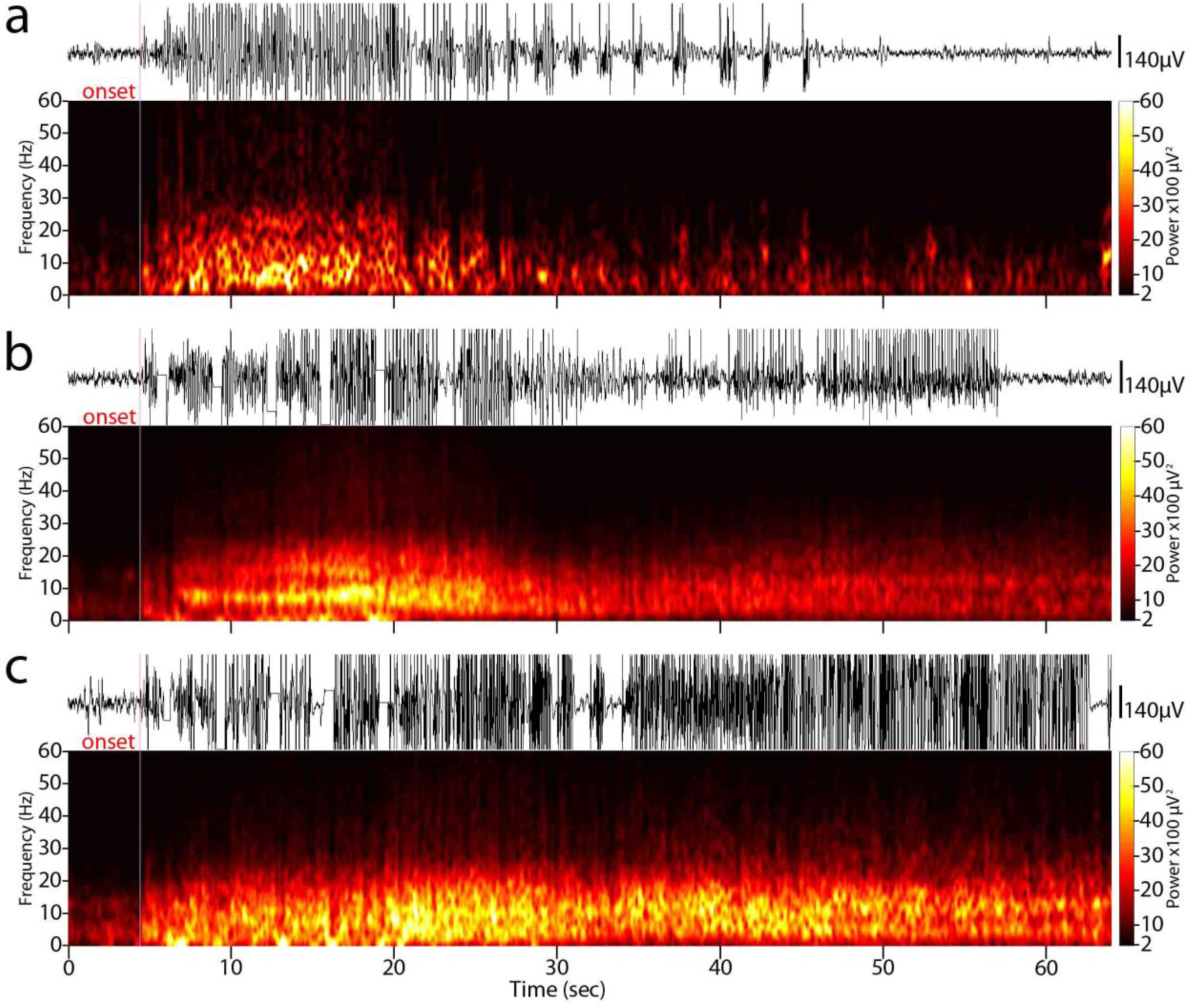
Indirect increase in mean electrographic seizure pattern duration. (a) Time-paired raw RNS data and averaged time-frequency power plots of patient 9’s electrographic seizure patterns from the baseline phase (n=2; channel 2; weeks 0-5). (b) During programming epoch weeks 5-14, a profound increase in mean electrographic seizure pattern duration was observed (n=17; channel 2). (c) Progressively, and up to programming epoch weeks 23- 37, the mean duration of electrographic seizure pattern exceeded the 60 s margin and reached a maximum of 79 s, representing a mean 132% increase (n=9; channel 2). Overall, for all patients, the mean onset of ictal duration modulation effects was 31 weeks following activation of stimulation.

## Abbreviations

RNS: responsive neuronal stimulator
ECoG: electrocorticography
PDMS: patient data management system
sEEG: stereotactic electroencephalography
EPSP: excitatory post-synaptic potential

